# Randomized gates eliminate bias in sort-seq assays

**DOI:** 10.1101/2022.02.17.480881

**Authors:** Brian L. Trippe, Buwei Huang, Erika A. DeBenedictis, Brian Coventry, Nicholas Bhattacharya, Kevin K. Yang, David Baker, Lorin Crawford

**Affiliations:** Massachusetts Institute of Technology, Cambridge, MA, USA; Microsoft Research New England, Cambridge, MA, USA; Institute for Protein Design, University of Washington, Seattle, WA, USA; Department of Biochemistry, University of Washington, Seattle, WA, USA; Department of Mathematics, University of California Berkeley, Berkeley, CA, USA; Howard Hughes Medical Institute, University of Washington, Seattle, WA, USA

## Abstract

Sort-seq assays are a staple of the biological engineering toolkit, allowing researchers to profile many groups of cells based on any characteristic that can be tied to fluorescence. However, current approaches, which segregate cells into bins deterministically based on their measured fluorescence, introduce systematic bias. We describe a surprising result: one can obtain unbiased estimates by incorporating randomness into sorting. We validate this approach in simulation and experimentally, and describe extensions for both estimating group level variances and for using multi-bin sorters.

---

Quantitative, multiplexed assays relying on fluorescence activated cell sorting (FACS) followed by high-throughput sequencing are critical to modern biology and molecular engineering because they enable construction of large scale datasets connecting sequence to function. For example, these “sort-seq” assays are widely used to profile the strength of protein-protein binding interactions via yeast display^4;5;11;17^. In particular one (i) synthesizes a *library* of 10^4^ to 10^5^ DNA sequences encoding proteins that may bind to a target of interest; (ii) transforms the library into yeast such that each putative binder is expressed on the surface of a population of cells; (iii) incubates cells with fluorescently labeled target protein; (iv) physically separates 10^6^ to 10^8^ cells based on binding affinity by FACS; and finally, (v) quantifies the prevalence, and thereby binding affinity, of each library member by high throughput sequencing. Due to biological and technical variability, there is a distribution over (log) fluorescence for each library sequence, and the challenge is to estimate the means of each of these distributions (Figure 1A-B). For example, for binding interactions, this mean fluorescence relates directly to biophysical quantities of interest including dissociation constants and binding energies^1;14;17^.

**Figure 1.**
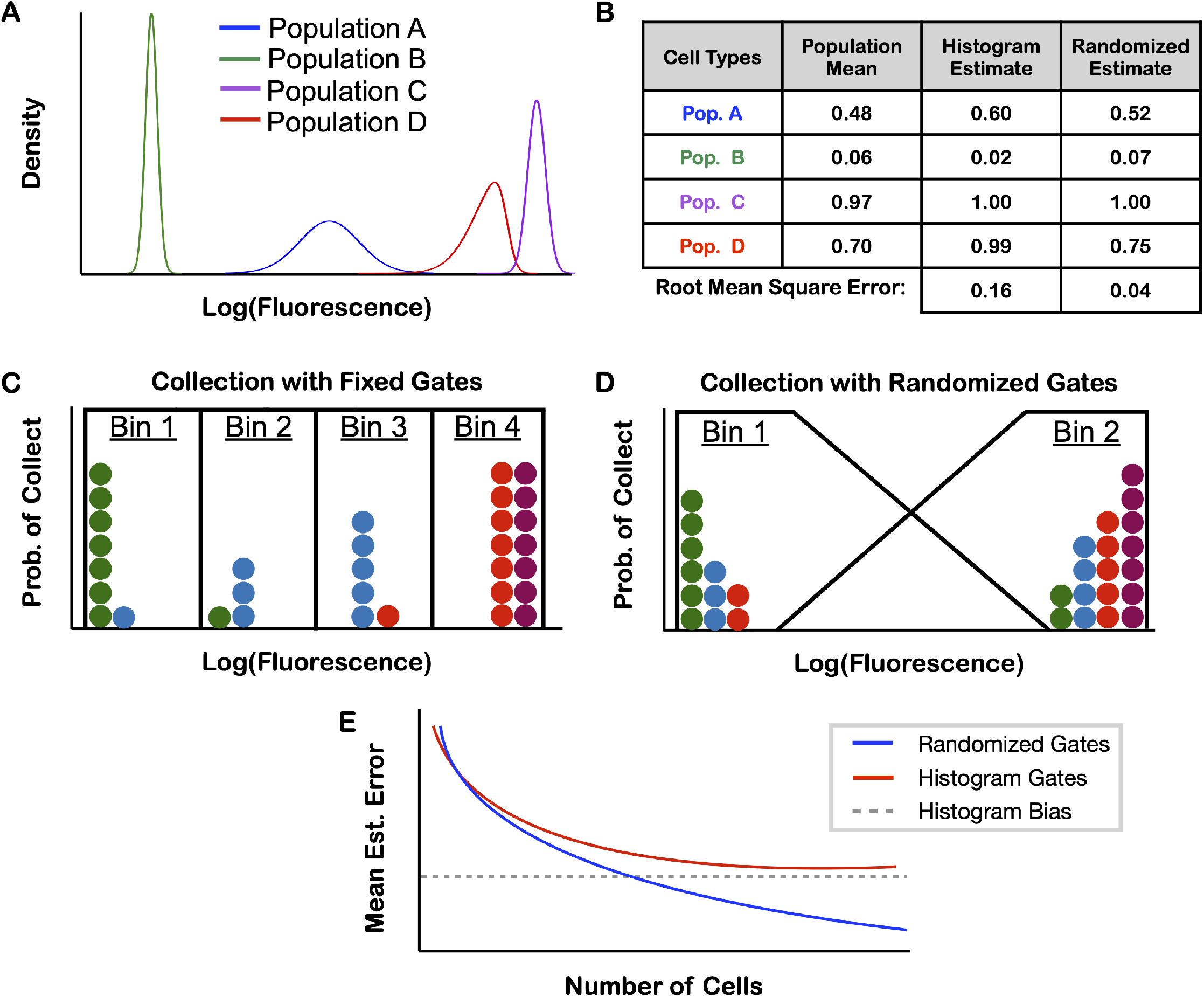
Schematic overview of randomized gates. **(A)** distributions of log fluorescence for different cell populations and **(B)** their hypothetical true and estimated means. **(C)** An example of histogram approach with deterministic collection into four bins and **(D)** an example of randomized collection approach with two bins. **(E)** Estimated means of the randomized gating scheme are more accurate than the histogram approach as the number of collected cells increases.

In previous work, cells are deterministically segregated into one or more collection tubes (referred to as “bins”) based on their measured fluorescences, and the mean fluorescence of each population is estimated from the histogram of observed sequence counts in each bin (Figure 1C). Peterman and Levine^13^ compare the error associated with different strategies for collecting and analyzing such data, and they show that average squared error is the sum of contributions from *bias* and *variance* (e.g., Hastie et al.^10^, Chapter 7.3). The variance arises from experimental noise and variability across cells, and it can be reduced by increasing the number of cells screened. The bias arises from the discretization of the space of log fluorescence into bins (Figure 1B-C); for example, narrow distributions can be sorted all into the same bin but have means as different as the bin width. Because even the most sophisticated FACS machines can sort cells into at most six bins, resolution is limited. This low resolution limits the value of sort-seq data in quantitative analyses, for instance, by prohibiting computation of precise binding energies. This challenge has spurred much work on how to effectively reduce histogram bias^1;8;13;14^. One common approach seeks to overcome the resolution limits of histograms by assuming fluorescence is log-normally distributed for each population and using maximum likelihood estimation to estimate moments^5;8;9;15^. However, on real data, this assumption is violated and the resulting estimates can have greater bias than the naive approach (Figure S1).

In this work, we show that the bias generated using histograms can be eliminated altogether by incorporating randomness into FACS collection strategies with as few as two bins (Figure 1D), thereby obtaining arbitrarily accurate estimates with many cells (Figure 1E). To do this, we take a statistical approach. We consider a population of cells that pass through a 2-bin sorter, each with log fluorescence *F* independently and identically distributed according to a density function *p*_*F*_. Our target of interest is the mean log fluorescence, ***μ***_*F*_ = ∫*fp*_*F*_ (*f*) d*f*. Let *B* denote the bin (either 1 or 2) into which a cell is collected, and let *Y*_1_ and *Y*_2_ be the counts of cells in bins 1 and 2 after sorting, respectively. In multiplexed sort-seq assays, we obtain *Y*_1_ and *Y*_2_ for thousands of populations, and our goal is to accurately estimate the mean of each population simultaneously.

For standard binning, a *gate* is chosen for each bin that defines the range of values *F* for which cells are collected into that bin; so, the bin *B* is deterministic once *F* is measured (e.g., as in Figure 1C). We instead consider *randomized gates* which define for each bin the probability of collecting a cell at each fluorescence (as in Figure 1D) and rely on pseudo-random numbers to determine the bin. For estimating population means, when the fluorescence measurements fall between lower and upper bounds *L* and *U*, one first sorts using randomized gates such that for any *f* on the interval [*L, U* ],

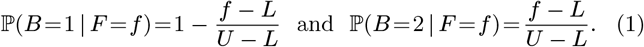

The counts are then combined into an empirical estimate of ***μ***_*F*_ as 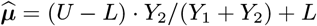

While one might expect introducing randomness to decrease precision by introducing additional noise, 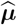 is directly informative to the mean fluorescence. In particular, 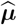 is an *unbiased* estimate of the true population mean in the sense that the average value we would expect for 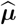 if we repeated the sort-seq experiment many times is equal to ***μ***_*F*_ (see Theorem 1 in Methods).

This unbiasedness theorem guarantees that, in contrast to the histogram approach, we can get arbitrarily accurate estimates by screening a larger numbers of cells (Figure 1E and Figure S2). More precisely, recalling that the mean squared error (MSE) is the sum of the bias squared and the variance^10^, unbiasedness implies that the error of 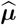 is dictated solely by its variance. Moreover, 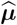 allows a transparent trade-off between the number of cells sorted per population and the precision of the estimates; notably, with as few as 400 cells, a 95% confidence interval for ***μ***_*F*_ will cover at most 10% of the range from *L* to *U* (Methods).

We used a simulation study to explore the implications of unbiasedness on estimation accuracy with the randomized gate approach relative to the standard histogram approach. In this study, we simulated fluorescence of 250 cells from log-normal distributions with different means and variances (Figure 2A). We then simulated sorting these cells based on their fluorescence either with four deterministic gates of equal width or with two randomized gates as dictated by Equation (1). For the deterministic gates, we constructed histograms and computed estimates of the mean fluorescence as the average of the bin centers weighted by the fraction of cells they contained; and for the randomized gates, we estimated the mean as 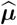. Figure 2B and C report the performance of these estimates in terms of MSE, along with their bias and variance components. As expected, the randomized gates approach has negligible bias except for broad distributions violating the conditions of our theorem (Methods).

**Figure 2.**
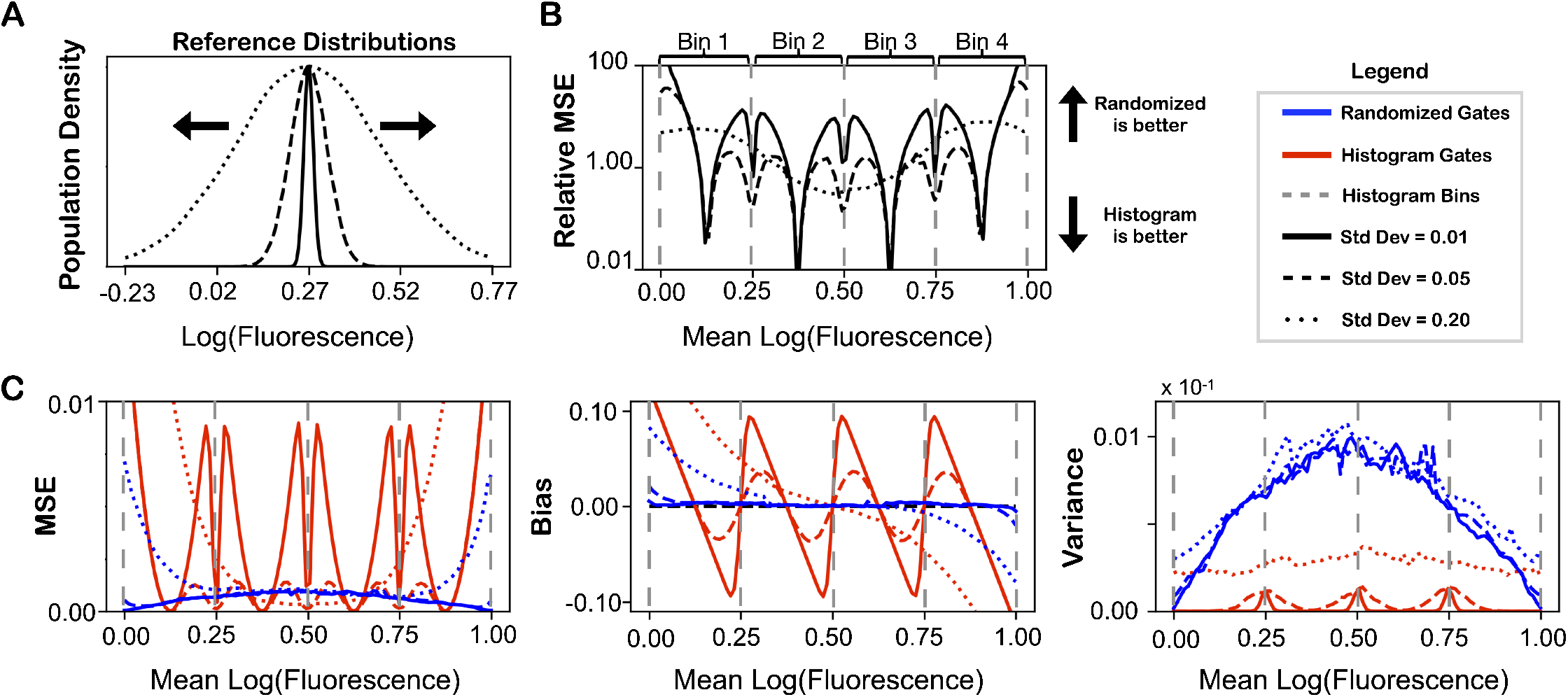
Simulation study reveals improved estimation properties obtained with randomized gates as compared to histograms. **(A)** Fluorescence values of cells are drawn independently from log-normal distributions with different scales. **(B)** The relative performances of estimates from histograms and randomized gates across a range of mean log fluorescences in terms of mean squared error (ratios greater than 1 reflect lower error with randomized gates and ratios below 1 reflect lower error with histograms). **(C)** The mean squared error (left) decomposed into bias (center) and variance (right) for both estimates. All points are the average across 200 replicates, each with *N* = 250 cells.

With even as few as 250 cells per population, the MSE of the histogram approach is dominated by bias. Accordingly, the unbiased randomized approach typically provides more accurate estimates. Notably, 250 cells is fewer than is the typical in sort-seq assays; with larger samples, more pronounced improvements are obtained (Figure S2). Because the histogram estimates are systematically biased toward bin centers, they can however be more accurate for narrow distributions with means near bin centers (Figure 2B).

We next tested our approach experimentally. Current FACS software does not support randomized gate programming, so we devised an experimental approximation in which we manually changed the gating threshold 20 times during sorting at regular intervals (Methods). We tested this procedure in the context of a binding assay using yeast display^3^. We synthesized DNA encoding four mini-protein binders to the SARS-COV-2 receptor binding domain (RBD) with a range of binding affinities^4^. While the value of this approach is greatest for highly multiplexed assays with many thousands of sequences, we chose this small number so that we could also test each binder easily in serial. We separately transformed and expressed each design in yeast and then incubated the populations with RBD. Both the target and binders were fluorescently labeled, and we considered the log ratio of target to binder fluorescence as an expression normalized proxy for binder strength^14^. We measured each sample on a Sony SH800 cell sorter separately, recording the binding signal for each binder (Figure 3A). We then pooled the samples together and sorted 1,000,000 cells, collecting 50,000 cells at each of the 20 thresholds (Methods).

**Figure 3.**
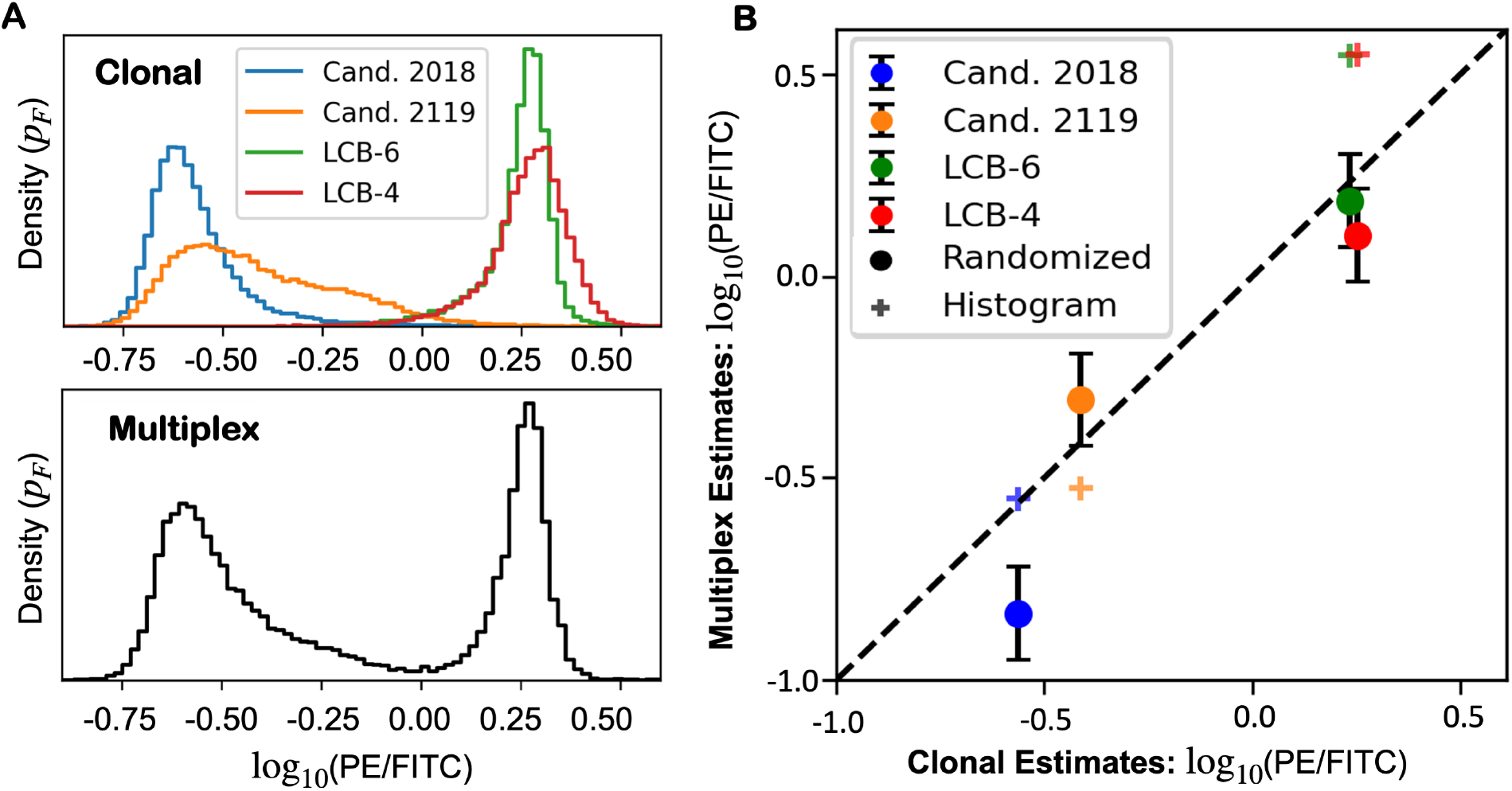
Agreement of binding signal of *de novo* designed binding proteins measured via yeast display in multiplex with ground truth values obtained in clonal yeast. **(A)** Distributions of samples measured clonally by flow cytometry, and distributions of pooled samples during sorting with a shifting gate boundary. **(B)** Agreement of clonal and multiplexed binding signal. The x-axis is measured by flow cytometry while the y-axis is a multiplexed measurement by next-generation sequencing. Error bars represent size of the steps used when shifting the threshold.

The multiplexed measurements largely recapitulate the ground-truth clonal measurements (Figure 3B), with the exception of design candidate 2018, for which the multiplexed estimate is below the clonal one. We suspect this is due to dissociation of some of the target protein in the time between the clonal and multiplexed measurements; kinetics experiments suggest dissociation occurs rapidly for this design^4^.

In the supplementary note, we additionally describe two extensions of this idea. First, because the differences in the variability of fluorescence across each population is often of interest (in addition to mean fluorescence), we show how to extend the approach to estimate the variance for each population. Second, we describe how to effectively take advantage of sorters that sort into more than two bins simultaneously to obtain more accurate estimates. We view these contributions as a starting point for future work of using randomness to obtain precise, multiplexed estimates.

We have shown how to obtain precise, multiplexed estimates in sort-seq experiments with a simple strategy that incorporates randomness. This mathematical technique allows better data to be collected using the same or less sophisticated hardware. While we have emphasized studies of binding affinity, we believe our strategy is applicable to a wider range of applications of sort-seq assays including studying transcriptional regulation^9;16^ and protein stability^15^, and building datasets for protein design^2^. Widespread implementation of randomized gates in FACS and community adoption of this strategy, will greatly simplify and improve sort-seq assays by eliminating a common bias in this ubiquitous assay. We believe this will allow FACS to play a more central role in screening settings, for construction of reliable datasets for machine learning models in bio-design applications, and for building datasets for quantitative models in biology more generally.

## Acknowledgements

We would like to thank Sarah Kate Nyquist for helpful conversations and suggestions. BLT would like to acknowledge support from the National Science Foundation Graduate Research Program. LC is supported by a David & Lucile Packard Fellowship for Science and Engineering. Any opinions, findings, and conclusions or recommendations expressed in this material are those of the author(s) and do not necessarily reflect the views of any of the funders.

## Methods

### Unbiasedness of estimates from randomized gates

The advantage of the randomized gates presented in Equation (1) is that the resulting counts in each bin (*Y*_1_ and *Y*_2_) may be combined as 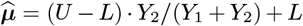 to estimate ***μ***_*F*_ without bias. We make this statement precise and present a theorem that guarantees when this is the case.

For an estimator 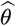 of a fixed estimand *θ*, the estimator’s *bias* is the expected value of its error 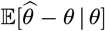 conditioned on that particular value of *θ*. An estimator is called *unbiased* if, regardless of the value of the estimand, the bias is equal to zero—that is, if 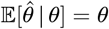 for every *θ*. Theorem 1 states that this property holds for 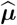.

#### Theorem 1

(Unbiasedness with randomized gates). *If the support of p*_*F*_ *is bounded between L and U, then* 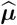 *is an unbiased estimator of mean fluorescence. That is*, 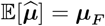.

*Proof*. We begin by rewriting the probability that a cell is collected into bin 2 to expose the connection between this quantity and ***μ***_*F*_ :

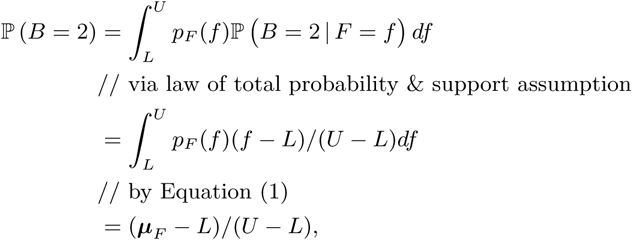

If *N* = *Y*_1_ + *Y*_2_ total cells are collected, then the count in the second bin is distributed as *Y*_2_|*N* ∼ Binomial ((***μ***_*F*_ *L*)*/*(*U L*),*N*) and has mean 𝔼[*Y*_2_] = *N* · (***μ***_*F*_–*L*)/(*U* − *L*). Accordingly, for any *N* total number of cells, 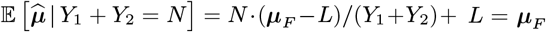. When *N* is random as well, then by the law of iterated expectation, 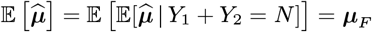 as desired. □

Notably, this theorem holds for any distribution *p*_*F*_ satisfying the support condition and does not require any parametric assumptions such as log-normality.

### Trade-off between number of cells sorted and precision of estimates

The relative simplicity of the estimate 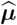 leads to a transparent trade-off between the precisio!n and scale of the experiment. Recalling that *Y*_2 |_*N* ∼ Binomial (***μ***_*F*_ − *L*)*/*(*U* − *L*),*N* the variance of 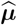 is

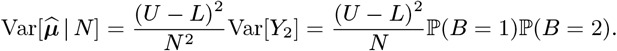

To construct a confidence interval for ***μ***_*F*_, we can therefore first approximate the standard error of 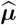 by 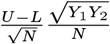, and appeal to approximate normality of the Binomial distribution for moderate to large *N* to report 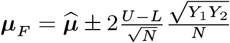 with 95% confidence. Because 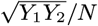 can be at most 1*/*2 (if *Y*_1_= *Y*_2_), the size of this interval is at most 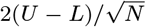. Therefore, to estimate ***μ***_*F*_ to within one tenth of the range with high confidence, at most *N* = 400 cells are needed, since in this case 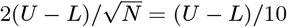.

For scale, commercial machines sort on the order of ten thousand cells per second, and typical assays sort tens of millions of cells divided amongst many populations. Thus, a library of one hundred thousand populations could be screened to high precision with on the order of 1 hour of sorting time.

### Simulation details

In the simulations depicted in Figure 2, we compare against the standard approach of using a histogram to estimate ***μ***_*F*_. Consider a *K* bin histogram. For each bin *k*, if the range of fluorescences collected is from lower bound *l*_*k*_ to upper bound *u*_*k*_, then ℙ (*B* = *k | F* = *f*) = **1**[*l*_*k*_ ≤*f < u*_*k*_]. The histogram estimate then corresponds to combining the resulting counts as

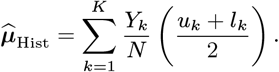

In order to use the unbiased estimator, both in simulation and in practice, we must slightly extend the randomized gate definition proposed in Equation (1). In particular, Theorem 1 assumes that the support of the fluorescence density *p*_*F*_ is bounded between *L* and *U* (i.e., that for *F ∼ p*_*F*_, ℙ [*L* ≤ *F* ≤ *U* ] = 1). In practice, this may not be the case. But, as previously stated, Equation (1) returns negative “probabilities” outside of this range. Therefore, we propose to “clip” the collection probabilities at the boundaries, and instead define

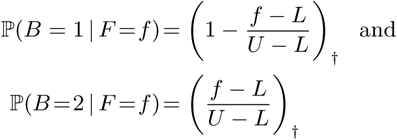

where † denotes clipping between zero and one such that, for a scalar *x*, (*x*)_†_ = max(min(*x*, 1), 0). This ensures that 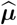 is well-defined, but gives up unbiasedness in situations where the support assumption of Theorem 1 is violated. This bias is apparent, for example, at the right and left sides of the left panel of Figure 2C.

### Experimental approximation of randomized gates with shifting thresholds

Because current FACS software does not support randomized gate programming, we devised an experimental approximation in which we manually changed the gating threshold 20 times during sorting at regular intervals. Specifically, we use a gate that collects all cells with fluorescence above a threshold into bin 2 and those below the threshold into bin 1, and we shift that threshold over the course of the collection from the lower limit *L* to the upper limit *U*. In theory, this approach exactly recovers Equation (1) in the limit that the threshold is shifted continuously from *L* to *U* at a constant rate. This is because for a cell with fluorescence *f* between *L* and *U*, the probability that it is collected into bin 2 is the fraction of the experimental time during which the threshold is below *f*, which is (*f* −*L*)*/*(*U* −*L*). This approximation does not, however, account for possible changes in the distribution, *p*_*F*_ over time. Such changes occur in binding assays, for example, when nontrivial labeled target protein dissociates over time. This challenge is a disadvantage of the approximation relative to randomized gates that could in theory be implemented into sorters.

### Yeast display and deep sequencing

EBY100 yeast cells expressing each of the four mini-protein binders were grown in C-Trp-Ura media. Binder protein expression was induced by replacing the growing buffer with SGCAA and incubating at 30° C for 24h^7^. The induced cells were labelled with 250 nM biotinylated receptor binding domain target protein, washed twice with PBSF (PBS+1% BSA), then labelled again with anti-c-Myc fluorescein isothiocyanate (FITC) and streptavidin-phycoerythrin (SAPE). The experiments were performed on a Sony SH800 cell sorter. 60,000 cells were recorded for each binder to reflect the individual distribution of baseline PE signal intensity. In the shifting gate experiment, a square area (`AreaTotal`) with side length (L) was pre-determined at the SH800 collection panel. The area was divided into 2 separate collection gates, `Gate1` and `Gate2` (corresponding to bin 1 and bin 2 in Equation (1)). `Gate2` was in an isosceles right triangle and started with a small area in the right-bottom corner of `AreaTotal` and `Gate1` took up the remaining. The yeast cells were run through the SH800 and each cell went into either the `Gate2` or `Gate1` collection tube if its log PE/FITC signal was in the range of `AreaTotal`. All other cells were discarded. After collecting 50,000 cells, the cell flow was paused, `Gate2` was shifted both leftwards and upwards for L/10 and cell flow continued. Because the proprietary software for operating the sorter allowed setting gate positions only through a point and click graphical user interface (rather than numerically), we measured out gate increments by pixel distance on the display using a ruler. The above shifting process repeated 19 times for a total of 20 collections. The cells collected in `Gate1` and `Gate2` were then grown, and 1 ×10^7^ cells from each gate were barcoded and the sequences for each cell were determined by Illumina next-generation sequencing^15^. The number of cells collected by each gate for each population was estimated from the proportion of sequencing reads attributed to each population and the number of cells collected into the gates.

Because the number of cells collected by each gate was not made directly available through the proprietary software, we estimated this from the raw exported data. In particular, we imported the data using the `FlowCal` python package^6^ and computationally implemented the gates and filters (including for forward and backward scatter).

### Sensitivity of maximum likelihood inference to non-normality of real data

Likelihood-based inference is a common strategy used with the intent to circumvent the resolution limitation of the histogram approach^5;8;9;15^. However, this approach can fail on real data. In particular, existing likelihood methods rely on the assumption that for each of the cell populations the fluorescence values are log normal distributed, log *F* ∼ 𝒩 (*μ, σ*^2^) where the mean log fluorescence *μ* = ***μ***_*F*_ is the target of inference and *σ*^2^ is the typically unknown variance of the population.

We evaluate performance of maximum likelihood inference in this situation with simulations using data sub-sampled from a flow cytometry dataset of binding signal of a computationally designed mini-protein binder to ActRII. Data were collected using yeast display as previously described except with the addition of a supplemental binding protein, protein A, the binding signal log(FITC*/*PE) was recorded for approximately one million cells. The distribution of this signal is highly non-Gaussian (Figure S1A).

We first compared the performance of the maximum likelihood approach (described in greater detail below) to the randomized approach on downsampled datasets with *N* = 250 cells with the same set-up described in Figure 2. As in the earlier simulations, the randomized approach provides improved MSE across most simulation conditions (Figure S1B). This improvement is again explained by estimation bias, which is mitigated by the randomized approach (Figure S1C). Though one might expect the benefit of maximum likelihood would appear for larger sample sizes (e.g., due to the asymptotic efficiency of maximum likelihood estimation in theory), this is not the case. In fact, due to the bias of maximum likelihood, the relative improvement of the randomized approach is larger at *N* = 1000 cells (Figure S1D). Moreover, Figure S1E demonstrates that the maximum likelihood approach does not empirically provide more accurate estimates even under correct specification (with fluorescences sampled as in Figure 2A).

### Maximum likelihood estimation

To estimate ***μ***_*F*_, likelihood-based approaches consider the counts in each of *K* bins (*Y*_1_, *Y*_2_,…, *Y*_*K*_), since the measured fluorescence values cannot be disambiguated when multiple populations are sorted in multiplex. These counts follow a multinomial distribution as

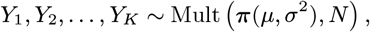

where 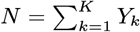 is the total number of cells sorted into any bin and *π*(*μ, σ*^2^)= (*π*_*1*,_*π*_*2*,_…, *π*_*K*_) are the normalized bin probabilities. In particular, if for each bin *k* the range of fluorescences collected is from lower bound *l*_*k*_ to upper bound *u*_*k*_, then

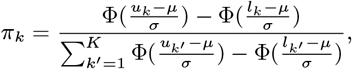

where Φ(·) is the cumulative density function of the standard normal. The log likelihood function is then

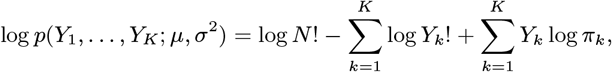

where the dependence of each *π*_*k*_ on *μ* and *σ*^2^ is left implicit. The maximum likelihood approach is to return *μ* that maximizes this expression,

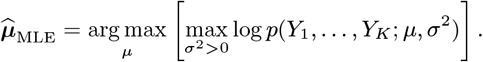

This optimization problem is not analytically tractable, and its constraints and non-convexity pose challenges for local, gradient-based optimizers. So we instead solve the optimization approximately with a grid search.

## Notes

### Competing Interest Statement

The authors have declared no competing interest.

