## Supplementary Note for "Randomized gates eliminate bias in sort-seq assays"

#### Estimating per-group variances with three randomized gates

In the main text, we considered estimating only population means. However, other distributional characteristics may also be of interest. For example, when fluorescence reports the expression level of a gene, the spread of the distribution $p_F$ reflects variability in transcription across cells. In this section, we describe an alternative set-up with three randomized gates that permit estimating the variance of $F$, which we denote by $\boldsymbol{\sigma}_F^2 = \int p_F(f)(f - \boldsymbol{\mu}_F)^2 df$. While our estimation approach is not unbiased for any finite number of cells (as was the case for $\widehat{\boldsymbol{\mu}}$), we demonstrate with theory and simulations that the estimates are accurate in assays with many cells.

Assume, for convenience, that the log fluorescences are between 0 and 1 (other ranges may be accommodated by rescaling accordingly). Let $B$ denote the bin into which a cell is collected (either 1, 2, or 3). We define the gates such that, conditional an any fluorescence $f$, $B$ is distributed as

$$B \mid F = f \sim \text{Categorical}\left[\left(f^2\right)_\dagger, \left(f - f^2\right)_\dagger, (1-f)_\dagger\right], \tag{S1}$$

where † again denotes clipping between 0 and 1. Figure S3A depicts these gates graphically.

We again let $Y_1, Y_2$, and $Y_3$ denote the number of cells collected into each bin, and let $N = Y_1 + Y_2 + Y_3$ be the total number of cells. We propose to estimate $\boldsymbol{\sigma}_F^2$ by $\widehat{\boldsymbol{\sigma}}^2 = N^{-1}Y_1 - \left[N^{-1}(Y_1 + Y_2)\right]^2$. Our next result justifies using this estimate. Specifically, we show that $\widehat{\boldsymbol{\sigma}}^2$ is *asymptotically consistent*, which means that the estimation error becomes arbitrarily small for a large enough number of cells.

**Theorem S1** (Asymptotic consistency of variance estimation)**.** *If $p_F$ is bounded between* 0 *and* 1 *then, in the limit of a large number of cells $N$, $\widehat{\boldsymbol{\sigma}}^2$ converges to $\boldsymbol{\sigma}_F^2$.*

*Proof.* From Equation (S1), $\mathbb{P}(B = 1 \mid F = f) = f^2$ for any $f$ between 0 and 1. By marginalizing over $f$, we obtain $\mathbb{P}(B = 1) = \int p_F(f)\mathbb{P}(B = 1 \mid F = f)df = \mathbb{E}[F^2]$. Thus, for any $N$, $Y_1 \mid N \sim \text{Binomial}(\mathbb{E}[F^2], N)$. By the law of large numbers, $N^{-1}Y_1 \to \mathbb{E}[F^2]$ as $N \to \infty$. Similarly, $N^{-1}Y_2 \to \mathbb{E}[F] - \mathbb{E}[F^2]$ and so, by the continuous mapping theorem, $[N^{-1}(Y_1 + Y_2)]^2 \to \mathbb{E}[F]^2$. Together, this implies that $\widehat{\boldsymbol{\sigma}}^2 \to \mathbb{E}[F^2] - \mathbb{E}[F]^2 = \boldsymbol{\sigma}_F^2$. □

The significance of Theorem S1 is that, similarly to Theorem 1 for means in the main text, it guarantees that one can accurately estimate variances in multiplex provided enough cells are sorted and sequenced. Notably, the randomized gates defined by Equation (S1) still permit unbiased estimation of means by $N^{-1}(Y_1 + Y_2)$.

**Estimating population variance in simulations.** In Figure S4, we evaluate the accuracy of $\widehat{\boldsymbol{\sigma}}^2$ in simulation across a range of distributions with different means and variances. As a result of Theorem S1, the error approaches zero as the number of cells increases. By contrast, the often large error of variance estimates obtained from histograms does improve with sample size.

#### Optimal unbiased multi-way sort-seq estimates

Thus far, we have shown only how to obtain estimates with an approach that sorts cells into one of two bins. However, many modern sorters allow up to 6-way sorting, placing each cell into one of 6 different bins. Utilizing a greater number of bins greatly improves the performance of histogram based estimates by lowering the discretization bias, and we want to take advantage of multi-way sorters with an unbiased randomized scheme. We describe one such procedure in the case of a 4-way sorter. We suspect that an extension of the procedure to a $K$-way sorting for arbitrary $K$ is straightforward. We prove that our proposal is optimal in the sense that, among a large class of possible estimation strategies, it has the smallest possible expected error for the worst case distribution $p_F$.

In order to approach the question of optimal randomized 4-way sorting rules, we consider a parameterized class of estimators for 4-way sorters. We separate the estimation strategies into two pieces: (i) the randomized gates and (ii) an analysis rule defining how the counts obtained are combined into a final estimate. We represent the gates by 4 functions $(\phi_1, \phi_1, \phi_2, \phi_4)$ such that for each gate $k$ a cell with fluorescence $f$ will be collected into bin $k$ with probability $\mathbb{P}(B = k \mid F = f) = \phi_k(f)$. As before, we let $Y_1, Y_2, Y_3$, and $Y_4$ denote the counts obtained in each bin. In contrast to the earlier cases, we do not restrict consideration to gates that collect all cells. For example, we allow $\sum_{k=1}^4 \phi_k(f) < 1$ for some $f$. This both allows for greater flexibility but also adds complication because the total number of cells sorted, which was key to the previous estimates, is no longer computable as $\sum_{k=1}^4 Y_k$. Instead, we assume that the expected number of cells sorted is known. In practice, this is easily obtained from the population proportions and the total number of cells sorted across all populations.

Next, we consider analysis rules that estimate $\boldsymbol{\mu}_F$ as an affine transformation of the counts, $\widehat{\boldsymbol{\mu}} = c_0 + \sum_1^4 c_k Y_k / \bar{N}$, where $c_0, \ldots, c_4$ are chosen constants that parameterize the analysis rule and $\bar{N}$ is the expected number of cells collected. While broader classes of analysis rules exist, we chose this class because it is (i) sufficiently expressive to contain useful estimates (e.g., choosing $\phi_1(f) = f, c_1 = 1$, and $c_k = 0$ for $k \neq 1$ provides an analogue of the 2-bin rule) and (ii) sufficiently constrained for us to reason about it precisely.

With this class, our task is then to characterize an effective set of gates $\boldsymbol{\phi} = (\phi_1, \ldots, \phi_4)$ and constants $\mathbf{c} = (c_0, \ldots, c_4)$ such that $\widehat{\boldsymbol{\mu}}$ remains unbiased and has the smallest possible error. Because we desire to have $\widehat{\boldsymbol{\mu}}$ perform well for any distribution $p_F$, we seek to find $\boldsymbol{\phi}$ and $\mathbf{c}$ that provide small error for the worst case $p_F$. For simplicity, we again assume the support of $p_F$ is bounded between 0 and 1.

This leads us to the constrained minimax optimization problem,

$$\begin{aligned}\boldsymbol{\phi}^*, \mathbf{c}^* = \underset{\boldsymbol{\phi},\mathbf{c}}{\arg\min}\max_{p_F}\mathbb{E}\left[(\widehat{\boldsymbol{\mu}} - \boldsymbol{\mu}_F)^2 \,|\, \boldsymbol{\phi}, \mathbf{c}\right], \\ \text{subject to (i) } \widehat{\boldsymbol{\mu}} \text{ is unbiased for } \boldsymbol{\mu}_F \text{ for } p_F \text{ supported on } [0,1], \\ \text{(ii) for each } f \text{ and } k,\ \phi_k(f) \geq 0, \text{ and} \\ \text{(iii) for each } f,\ \sum_{k=1}^{4}\phi_k(f) \leq 1.\end{aligned} \tag{S2}$$

The three constraints above correspond to (i) unbiasedness of the estimator, (ii) that gates cannot collect cells with negative probability, and (iii) that the sum of the probabilities of collecting any cell may not exceed 1.

Equation (S2) is a challenging, infinite-dimensional, and non-convex optimization problem. Nevertheless, we are able to characterize a solution: $\phi_1^*(f) = \mathbf{1}_{(-\infty,\frac{1}{4}]}(f)\left[2(\frac{1}{2}-f)\right]_\dagger$; $\phi_2^*(f) = \mathbf{1}_{(\frac{1}{4},\frac{1}{2})}(f)\left[4(\frac{1}{2}-f)\right]$; $\phi_3^*(f) = \mathbf{1}_{(\frac{1}{2},\frac{3}{4})}(f)\left[4(f-\frac{1}{2})\right]$; $\phi_4^*(f) = \mathbf{1}_{[\frac{3}{4},\infty)}(f)\left[2(f-\frac{1}{2})\right]_\dagger$; and $[c_0^*, c_1^*, c_2^*, c_3^*, c_4^*] = \left[\frac{1}{2}, -\frac{1}{2}, -\frac{1}{4}, \frac{1}{4}, \frac{1}{2}\right]$, where $\mathbf{1}_A$ is the indicator function for membership in the set $A$. Figure S3B depicts these gates graphically. Our third theorem formally states the optimality of this solution.

**Theorem S2** (Minimaxity of $\boldsymbol{\phi}$ and $\mathbf{c}^*$)**.** *Suppose the number of cells sorted is distributed as* $N \sim \text{Poisson}(\bar{N})$*, then the gates* $\boldsymbol{\phi}^*$ *and constants* $\mathbf{c}^*$ *provide minimax optimal expected squared error among the class of unbiased 4-way sorting rules described above for densities* $p_F$ *supported on* $[0,1]$.

Theorem S2 provides two useful guarantees about $\boldsymbol{\phi}^*$ and $\mathbf{c}^*$. First, it assures that the estimates remain unbiased as so will perform well in the very high-throughput regime. And second, it provides that the worst case performance of this approach is as good as is possible for estimators within this large class.

We provide a proof of the theorem in the next section. In brief, we first solve the inner optimization problem analytically using $L_1$ - $L_\infty$ duality. This provides a more tractable optimization problem, subject to linear constraints, whose objective is the $L_\infty$ norm of an affine combination the $\phi_k$'s. The result is then obtained by showing that any $(\boldsymbol{\phi}, \mathbf{c})$ satisfying the conditions of Equation (S2) has loss at least as large as $(\boldsymbol{\phi}^*, \mathbf{c}^*)$.

We note that Theorem S2 does not, however, guarantee that the solution to Equation (S2) provided above is unique. In fact, a subspace of solutions with equal maximum expected error exists.

### Proof of Theorem S2

*Proof.* To prove Theorem S2 we begin by rewriting the minimax problem with a simplified objective and more explicit constraints. We then derive a lower bound on the minimax objective and show that the proposed solution $(\boldsymbol{\phi}^*, \mathbf{c}^*)$ meets this bound. Since this solution meets the constraints of the problem by construction, it is therefore minimax optimal. For notational convenience, let $\mathcal{P}$ denote the set of all densities $p_F$ supported on $[0,1]$. Notably, we recognize that $\mathcal{P}$ is the set of positive functions in $\mathrm{L}_1[0,1]$ with unit norm; that is, for any $p_F$ in $\mathcal{P}$, we have $\|p_F\|_1 = \int_0^1 p_F(f)df = 1$.

We begin by reformulating the minimax problem. First, Lemma S1 proves that the unbiasedness condition is equivalent to that for almost every $f \in [0,1]$, $c_0 + \sum_{k=1}^K c_k\phi_k(f) = f$, where $K$ is the number of bins ($K = 4$ in Equation (S2)). Next, Lemma S2 proves that the objective of the inner maximization problem may be written as an inner product in the function space $\mathrm{L}_2[0,1]$. Lemma S3 gives the solution to this maximization problem in closed form. Together, these allow us to rewrite the minimax problem as

$$\begin{aligned}\boldsymbol{\phi}^*, \mathbf{c}^* = \underset{\boldsymbol{\phi}\,\in\,\mathrm{L}_\infty^K[0,1],\,\mathbf{c}\,\in\,\mathbb{R}^{K+1}}{\arg\min}\left\|\sum_{k=1}^{K} c_k^2\phi_k\right\|_\infty \\ \text{subject to (i) for almost every } f \in [0,1],\ c_0 + \sum_{k=1}^{K} c_k\phi_k(f) = f, \\ \text{(ii) for each } f \in [0,1] \text{ and } k,\ \phi_k(f) \geq 0, \text{ and} \\ \text{(iii) for each } f \in [0,1],\ \sum_{k=1}^{K}\phi_k(f) \leq 1,\end{aligned} \tag{S3}$$

where $\mathrm{L}_\infty^K[0,1]$ is the set whose elements each consist of $K$ functions in $\mathrm{L}_\infty^K[0,1]$. Next, Lemma S4 shows that to solve Equation (S3) it suffices to consider solutions satisfying the additional condition that $\|\phi_k\|_\infty = 1$ for each $k$. Specifically, this lemma states that any $(\boldsymbol{\phi}, \mathbf{c})$ satisfying the conditions of Equation (S3), but violating this additional condition, may be modified to satisfy the condition without increasing the loss.

Lemma S5 provides a lower bound on the objective for all $(\boldsymbol{\phi}, \mathbf{c})$ satisfying all four conditions. In particular, every feasible $(\boldsymbol{\phi}, \mathbf{c})$ has $\|\sum_{k=1}^K c_k^2\phi_k\|_\infty \geq 1/4$.

Lastly, we conclude the proof by noting that the proposed solution $(\boldsymbol{\phi}^*, \mathbf{c}^*)$ satisfies the conditions of Equation (S3) and attains this minimum loss. The conditions are satisfied by construction. Because the values $f \in [0,1]$ for which each $\phi_k^*(f)$ is positive do not overlap and because for each $k$ $\|\phi_k^*\|_\infty = 1$, the loss simplifies as $\|\sum_{k=1}^K c_k^{*2}\phi_k^*\|_\infty = \max_{1\leq k\leq K} c_k^{*2}\|\phi_k^*\|_\infty = \max_{1\leq k\leq K} c_k^{*2} = 1/4$. Therefore, $(\boldsymbol{\phi}^*, \mathbf{c}^*)$ minimizes Equation (S3) and $\widehat{\boldsymbol{\mu}}$ is minimax optimal. ☐

**Lemma S1.** *The unbiasedness condition on* $(\boldsymbol{\phi}, \mathbf{c})$*,*

$$\text{for any } p_F \in \mathcal{P},\ \mathbb{E}_{p_F}[\widehat{\boldsymbol{\mu}} \mid \boldsymbol{\phi}, \mathbf{c}] = \boldsymbol{\mu}_F, \tag{S4}$$

*is equivalent to*

$$\text{for almost every } f \in [0,1],\ c_0 + \sum_{k=1}^{K} c_k \phi_k(f) = f, \tag{S5}$$

*where we mean 'almost every' in the measure theoretic sense that Equation* (S5) *holds on* $[0,1]$ *except on a set of Lebesgue measure zero.*

*Proof.* We prove equivalance be showing that each Equation (S4) and Equation (S5) imply one another. First assume that Equation (S5) holds. Then, for any $p_F \in \mathcal{P}$, we may write

$$\begin{aligned}
\mathbb{E}_{p_F}[\widehat{\boldsymbol{\mu}} \mid \boldsymbol{\phi}, \mathbf{c}] &= \mathbb{E}_{N\sim\mathrm{Pois}(\bar{N})}\left[\mathbb{E}_{p_F}[c_0 + N^{-1}\sum_{k=1}^{K} c_k Y_k \mid \boldsymbol{\phi}, \mathbf{c}, N]\right] \\
&\text{// by the law of iterated expectation} \\
&= \mathbb{E}_{p_F}\left[c_0 + \sum_{k=1}^{K} c_k Y_k \mid \boldsymbol{\phi}, \mathbf{c}, N = 1\right] \\
&\text{// since multinomial expectations scale linearly in } N \\
&= \mathbb{E}_{p_F}\left[\mathbb{E}[c_0 + \sum_{k=1}^{K} c_k Y_k \mid \boldsymbol{\phi}, \mathbf{c}, N = 1, F]\right] \\
&\text{// by the law of iterated expectation} \\
&= \mathbb{E}_{p_F}\left[c_0 + \sum_{k=1}^{K} c_k \phi_k(F)\right] \\
&= \mathbb{E}_{p_F}[F] \\
&= \boldsymbol{\mu}_F.
\end{aligned}$$

It remains to show that Equation (S4) implies Equation (S5). We assume for contradiction that $(\boldsymbol{\phi}, \mathbf{c})$ does not satisfy Equation (S5) and will show that $\widehat{\boldsymbol{\mu}}$ can be biased. By assumption, there is a subset $A \subset [0,1]$ with positive Lebesgue measure such that for all $f \in A$, $c_0 + \sum c_k \phi_k(f) \neq f$. Let $A_+$ and $A_-$ be the subsets of $A$ on which $c_0 + \sum c_k \phi_k(f)$ exceeds and falls below $f$, respectively. At least one of $A_+$ and $A_-$ has positive measure; here, we assume the $A_+$ does (the other case follows symmetrically). Next, consider the density $p_F(f) = \lambda(A_+)^{-1}\mathbf{1}[f \in A_+]$, where $\lambda(A_+)$ is the Lebesgue measure of $A_+$. Then

$$\begin{aligned}
\mathbb{E}_{p_F}[\widehat{\boldsymbol{\mu}} \mid \boldsymbol{\phi}, \mathbf{c}] - \boldsymbol{\mu}_F &= \mathbb{E}_{p_F}\left[\mathbb{E}[c_0 + \bar{N}^{-1}\sum_{k=1}^{K} c_k Y_k \mid F] - F\right] \\
&= \lambda(A_+)^{-1}\int_{A_+} (c_0 + \sum_{k=1}^{K} c_k \phi_k(f)) - f df \\
&> 0.
\end{aligned}$$

Therefore, violations of this condition imply $\widehat{\boldsymbol{\mu}}$ is biased for some $p_F$ and Equation (S4) does not hold. The two conditions are therefore equivalent. □

**Lemma S2.** *Consider* $(\boldsymbol{\phi}, \mathbf{c})$ *satisfying the conditions of Equation* (S3)*. Then, for any* $p_F \in \mathcal{P}$*,*

$$\mathbb{E}_{p_F}[(\widehat{\boldsymbol{\mu}} - \boldsymbol{\mu}_F)^2 \mid \boldsymbol{\phi}, \mathbf{c}] = \bar{N}^{-1}\left\langle \sum_{k=1}^{K} c_k^2 \phi_k,\ p_F \right\rangle,$$

*where* $\langle \cdot, \cdot \rangle$ *denotes the inner product on* $\mathrm{L}_2$*. That is, for functions* $f, g \in \mathrm{L}_2[0,1]$, $\langle f, g \rangle = \int_0^1 f(x)g(x)dx$*.*

*Proof.* We begin by recognizing that since (by Lemma S1) condition (i) provides that $\widehat{\boldsymbol{\mu}}$ is unbiased, the expected squared error of $\widehat{\boldsymbol{\mu}}$ is equal to its variance:

$$\mathbb{E}_{p_F}[(\widehat{\boldsymbol{\mu}} - \boldsymbol{\mu}_F)^2 \mid \boldsymbol{\phi}, \mathbf{c}] = \mathrm{Var}[\widehat{\boldsymbol{\mu}} \mid \boldsymbol{\phi}, \mathbf{c}].$$

We next derive a reduced expression for the variance. To do this, we observe that the counts $Y_k$ in each bin are independently Poisson distributed as $Y_k \overset{indep.}{\sim}$ Pois $\left(\bar{N}\mathbb{E}[Y_k \mid \boldsymbol{\phi}, \mathbf{c}, N=1]\right)$. Therefore the variance simplifies as

$$\begin{aligned}
\text{Var}[\widehat{\boldsymbol{\mu}} \mid \boldsymbol{\phi}, \mathbf{c}] &= \text{Var}\left[c_0 + \bar{N}^{-1}\sum_{k=1}^{K} c_k Y_K \mid \boldsymbol{\phi}, \mathbf{c}\right] \\
&= \bar{N}^{-2}\text{Var}\left[\sum_{k=1}^{K} c_k Y_K \mid \boldsymbol{\phi}, \mathbf{c}\right] \\
&= \bar{N}^{-2}\sum_{k=1}^{K} c_k^2 \text{Var}\left[Y_K \mid \boldsymbol{\phi}, \mathbf{c}\right]
\end{aligned}$$

This independence is a general property of Poisson-multinomial models; that is, for any multinomial probabilities $\theta = (\theta_1, \ldots, \theta_K)$, if $N \sim \text{Pois}(\bar{N})$ and $Y_1, \ldots, Y_K \mid N \sim \text{Mult}(\theta, N)$, then each $Y_k \overset{indep.}{\sim} \text{Pois}(\bar{N}\theta_k)$.

Next, again because each $Y_k$ is Poisson distributed, the variance of each $Y_k$ is equal to its mean. Thus, $\text{Var}[Y_k \mid \boldsymbol{\phi}, \mathbf{c}] = \bar{N}\int_0^1 p_F(f)\phi_k(f)df$. We obtain the result by recognizing that $\int_0^1 p_F(f)\phi_k(f)df = \langle p_F, \phi_k\rangle$, where $\langle\cdot,\cdot\rangle$ is the inner product on $\text{L}_2[0,1]$, and then simplifying our earlier expression as

$$\begin{aligned}
\text{Var}[\widehat{\boldsymbol{\mu}} \mid \boldsymbol{\phi}, \mathbf{c}] &= \bar{N}^{-2}\sum_{k=1}^{K} c_k^2 \text{Var}[Y_K \mid \boldsymbol{\phi}, \mathbf{c}] \\
&= \bar{N}^{-1}\sum_{k=1}^{K} c_k^2 \langle \phi_k, p_F\rangle \\
&= \bar{N}^{-1}\langle \sum_{k=1}^{K} c_k^2 \phi_k, p_F\rangle.
\end{aligned}$$

□

**Lemma S3.** *Consider $f \in \text{L}_\infty[0,1]$ satisfying $f(t)$ for all $t$ in $[0,1]$. Then*

$$\sup_{x\in\mathcal{P}} \langle x, f\rangle = \|f\|_\infty,$$

*where $\|\cdot\|_\infty$ denotes the $\text{L}_\infty$ norm of a function (i.e., $\|g\|_\infty = \text{ess}\sup_t |g(t)|$, where ess sup denotes the essential supremum).*

*Proof.* Consider first the optimization problem on a larger domain, the unit sphere in $\text{L}_1[0,1]$. Duality of the function spaces $\text{L}_1$ and $\text{L}_\infty$ ($\text{L}_\infty$ is the dual space of $\text{L}_1$), provides that

$$\sup_{x\in \text{L}_1,\, \|x\|\le 1} \langle x, f\rangle = \|f\|_\infty. \tag{S6}$$

See for example Luenberger [12] (Chapter 5.8, Theorem 2; and Chapter 5.9, Example 2) taking the subspace ("$M$" in their notation) to be all of $\text{L}_1$. Since $\mathcal{P}$ is a subset of this unit sphere, Equation (S6) provides an upper bound on the objective. To see that this upper bound is attained, we can consider a sequence of level sets $A_n = \{t \in [0,1] \mid f(t) \ge \|f\|_\infty - n^{-1}\}$ and densities $p_F^{(n)}$ in $\mathcal{P}$ with positive density only on these level sets. The level sets have positive Lebesgue measure for any $f \in L_\infty[0,1]$, therefore, guaranteeing the existence of the sequence $\{p_F^{(n)}\}_{n=1}^N$. By construction, $\lim_{n\to\infty}\langle p_F^{(n)}, f\rangle = \|f\|_\infty$, and thus $\sup_{x\in\mathcal{P}}\langle x, f\rangle = \|f\|_\infty$. □

**Lemma S4.** *For any $(\boldsymbol{\phi}, \mathbf{c})$ satisfying the conditions of Equation (S3), there exists $(\boldsymbol{\phi}', \mathbf{c}')$ also satisfying the conditions of Equation (S3) with $\|\phi_k'\|_\infty = 1$ for each $k$ and $\|\sum_{k=1}^K c_k'^2\phi_k'\|_\infty \le \|\sum_{k=1}^K c_k^2\phi_k\|_\infty$.*

*Proof.* If $(\boldsymbol{\phi}, \mathbf{c})$ already satisfies the condition, let $(\boldsymbol{\phi}', \mathbf{c}') = (\boldsymbol{\phi}, \mathbf{c})$. Otherwise, consider each bin $k$ such that $\|\phi_k\|_\infty = d_k < 1$. Next define $(\boldsymbol{\phi}', \mathbf{c}')$ with each $\phi_k' = \phi_k/d_k$ and $c_k' = d_k c_k$.

We next consider the difference in loss for the unadjusted solution $(\boldsymbol{\phi}, \mathbf{c})$, as compared to the adjusted solution $(\boldsymbol{\phi}', \mathbf{c}')$. For any $p_F$,

$$\begin{aligned}
\bar{N}^{-1}\left\langle \sum_{k=1}^{K} c_k^2\phi_k, p_F\right\rangle - \bar{N}^{-1}\left\langle \sum_{k=1}^{K} c_k'^2\phi_k', p_F\right\rangle &= \bar{N}^{-1}\sum_{k=1}^{K}\left\langle c_k^2\phi_k - c_k'^2\phi_k', p_F\right\rangle \\
&= \bar{N}^{-1}\sum_{k=1}^{K}\left\langle c_k^2\phi_k - \frac{d_k^2}{d_k}c_k^2\phi_k, p_F\right\rangle \\
&= \bar{N}^{-1}\sum_{k=1}^{K}(1-d_k)\langle \phi_k, p_F\rangle \\
&\ge 0
\end{aligned}$$

and therefore we see that $(\boldsymbol{\phi}', \mathbf{c}')$ can have only smaller loss than $(\boldsymbol{\phi}, \mathbf{c})$. □

**Lemma S5.** *Suppose* $(\boldsymbol{\phi}, \mathbf{c})$ *satisfies the conditions of Equation* (S3) *and that* $\|\phi_k\|_\infty = 1$ *for each* $k$. *Then* $\|\sum_{k=1}^K c_k^2 \phi_k\|_\infty \geq 1/4$.

*Proof.* To prove the lemma, we consider two cases ($c_0 \leq 1/2$ or $c_0 > 1/2$). In the first case, we assume $c_0 \leq 1/2$. Then by condition (i) of Equation (S3), for any $\epsilon > 0$, there is some $f > 1 - \epsilon$ such that $c_0 + \sum_{k=1}^K c_k \phi_k(f) = f \geq 1 - \epsilon$. Therefore, $\sum_{k=1}^K c_k \phi_k(f) \geq 1 - c_0 - \epsilon \geq \frac{1}{2} - \epsilon$. Next note that $\sum_{k=1}^K c_k \phi_k(f) \leq (\max_{1 \leq k \leq K} c_k) \sum_{k=1}^K \phi_k(f) \leq \max_{1 \leq k \leq K} c_k$. Together, this implies that there is some $k$ such that $c_k \geq \frac{1}{2} - \epsilon$. Since by assumption each $\phi_k$ satisfies $\|\phi_k\|_\infty = 1$, the optimal objective is at least $\frac{1}{4}$, since $\|\sum_{k=1}^K c_k^2 \phi_k\|_\infty \geq \|c_k^2 \phi_k\|_\infty \geq (\frac{1}{2} - \epsilon)^2 > \frac{1}{4} - 2\epsilon$. Because $\epsilon$ is arbitrary, we see that $\|\sum_{k=1}^K c_k^2 \phi_k\|_\infty \geq 1/4$.

The second case ($c_0 > 1/2$) follows similarly, except that as the intermediate step instead of $c_k \geq \frac{1}{2} - \epsilon$ we obtain $c_k \leq -1/2$. Since these two cases are exhaustive, the result follows. □

### Supplementary Figures

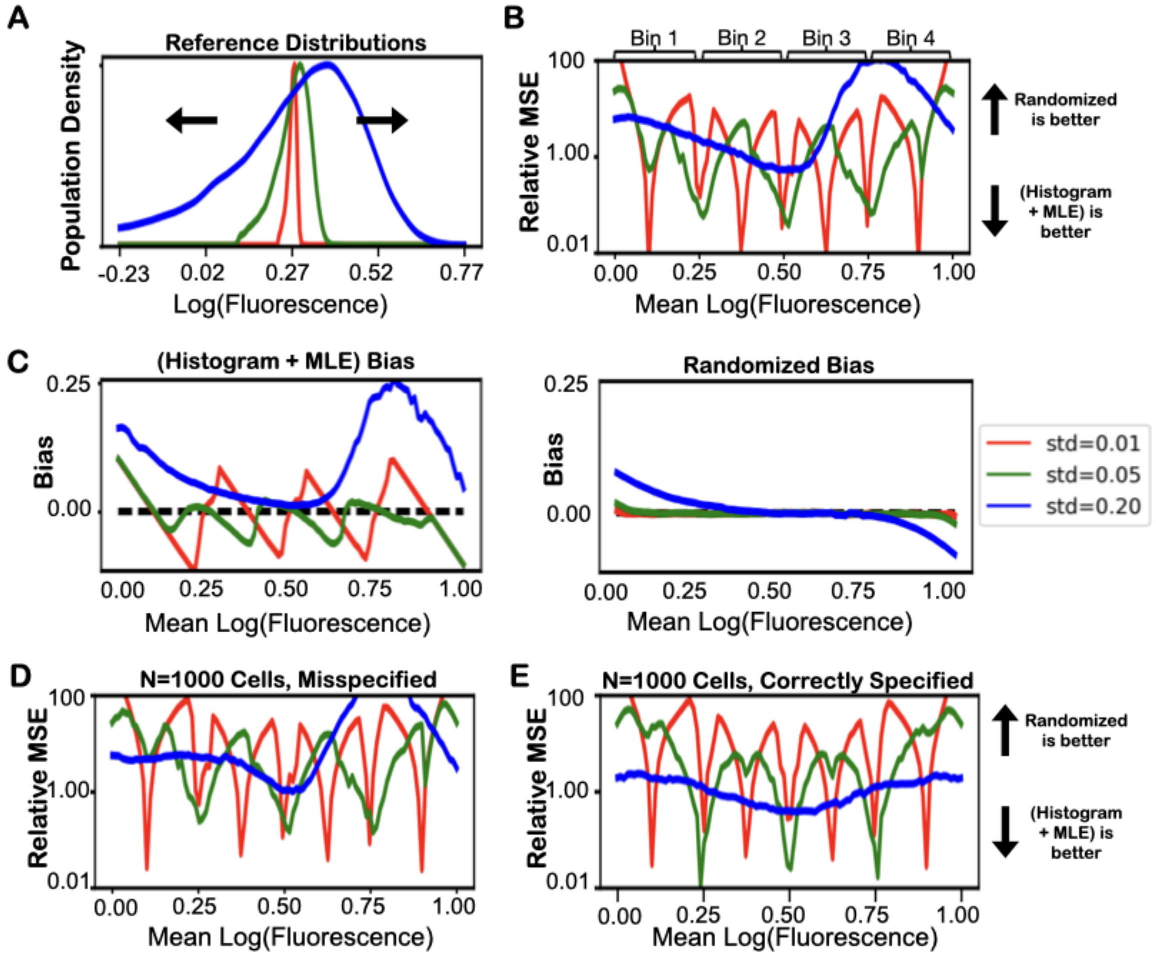


**Figure S1. Maximum likelihood estimation of mean fluorescence assuming log normality is non-robust on real data.** **(A)** Fluorescence values are drawn independently from empirical distributions of binding signal of mini-protein binders to ActRII obtained by flow cytometry and rescaled to have an range of different standard deviations (0.01 in red, 0.05 in green, and 0.20 in blue, respectively). **(B)** The ratio of the MSE of estimates from the maximum likelihood approach on histogram counts to unbiased estimates from randomized gates. **(C)** The estimation bias for the maximum likelihood estimates with the histogram counts (left) versus the randomized gates (right). All points in panels **(B)** and **(C)** are the average across 200 replicates, each with $N = 250$ cells. **(D)** The ratio of the MSE of the estimates is not substantially different even with a greater number ($N = 1000$) of cells or **(E)** under correct specification with log normal distributed observations.

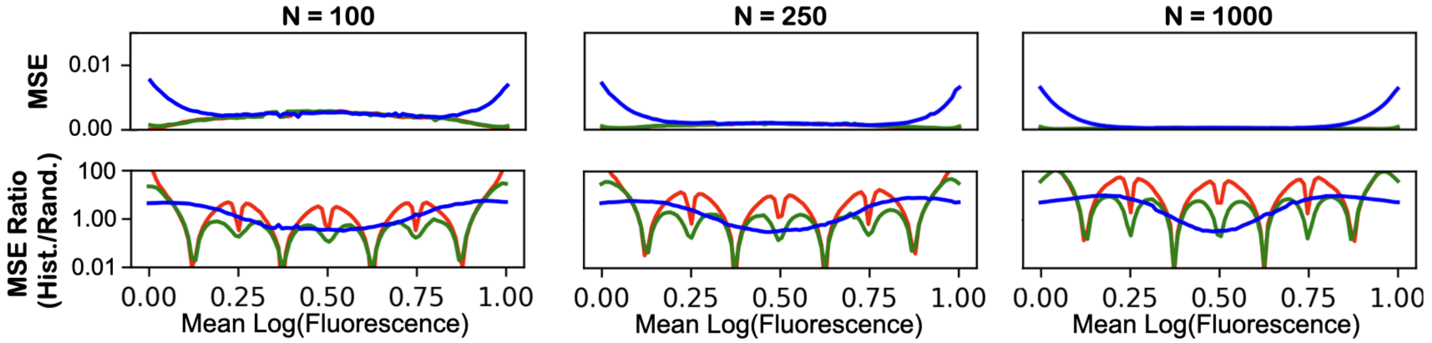


**Figure S2. The error of estimates with randomized gates decreases with increasing sample size.** Here, we consider $N = 100$, 250, and 1000 total cells, respectively. The bottom row shows the relative performances of estimates from histograms and randomized gates across a range of mean log fluorescences in terms of mean squared error (ratios greater than 1 reflect lower error with randomized gates and ratios below 1 reflect lower error with histograms).

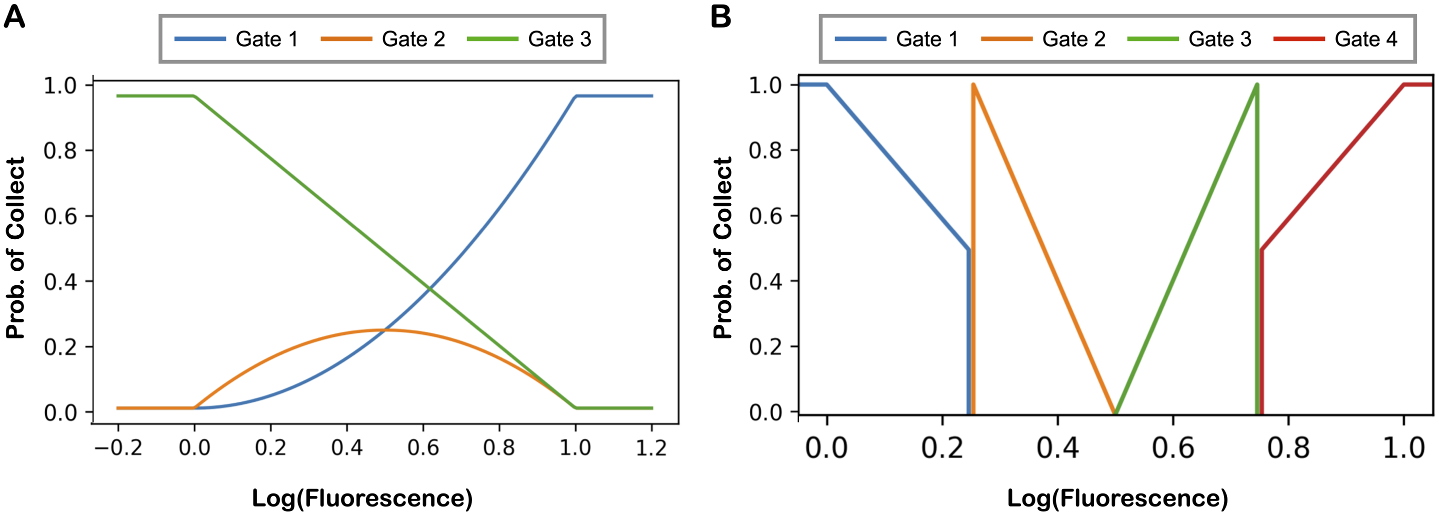


**Figure S3.** Randomized gates for **(A)** estimating population variances as defined by Equation (S1) and **(B)** optimally estimating population means with 4-way sorting.

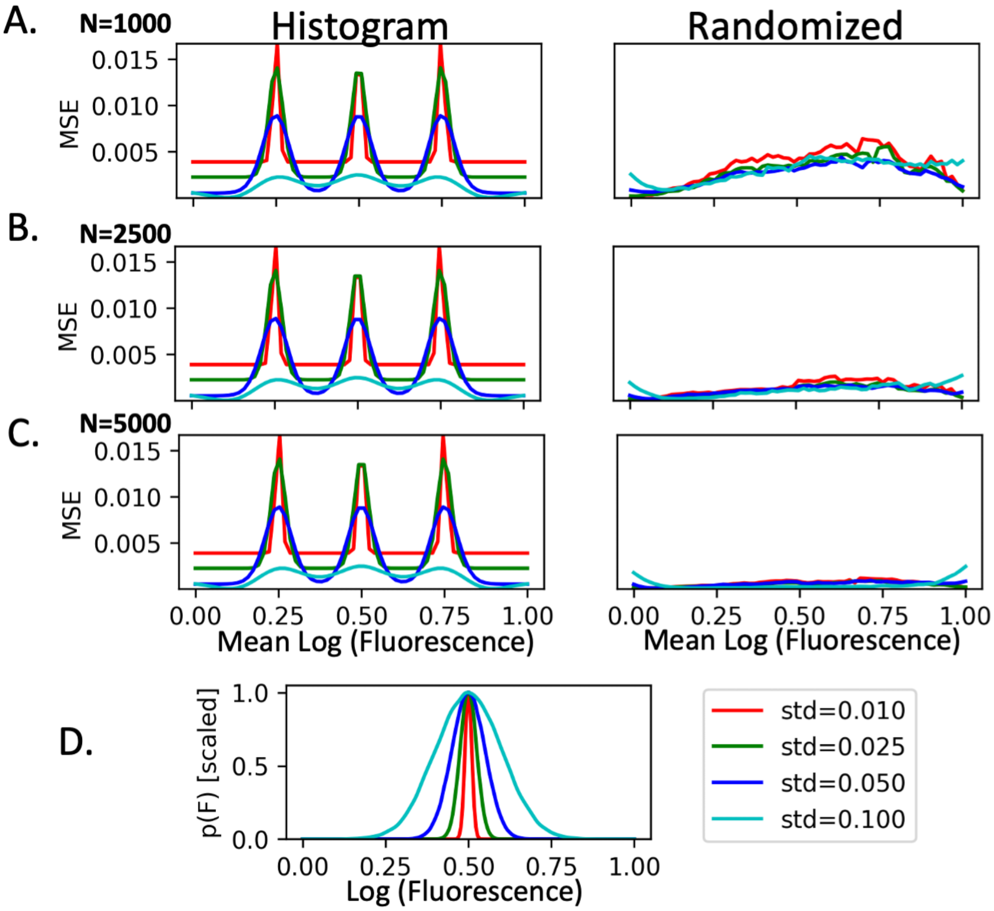


**Figure S4. Average squared error of standard deviation estimates of various $p_F$ at different sample sizes.** Here, we consider **(A)** $N = 1000$, **(B)** $N = 5000$, and **(C)** $N = 25000$ total cells. Depicted are the performances of (left) histogram based estimates and (right) the randomized estimate $\sqrt{\widehat{\boldsymbol{\sigma}}^2}$. In panel **(D)**, we show the distributions from which we simulated the data.
